# Random and natural non-coding RNA have similar structural motif patterns but can be distinguished by bulge, loop, and bond counts

**DOI:** 10.1101/2022.09.01.506257

**Authors:** Fatme Ghaddar, Kamaludin Dingle

## Abstract

An important question in evolutionary biology is whether and in what ways genotype-phenotype (GP) map biases can influence evolutionary trajectories. Untangling the relative roles of natural selection and biases (and other factors) in shaping phenotypes can be difficult. Because RNA secondary structure (SS) can be analysed in detail mathematically and computationally, is biologically relevant, and a wealth of bioinformatic data is available, it offers a good model system for studying the role of bias. For quite short RNA (length *L* ≤ 126), it has recently been shown that natural and random RNA are structurally very similar, suggesting that bias strongly constrains evolutionary dynamics. Here we extend these results with emphasis on much larger RNA with length up to 3000 nucleotides. By examining both abstract shapes and structural motif frequencies (ie the numbers of helices, bonds, bulges, junctions, and loops), we find that large natural and random structures are also very similar, especially when contrasted to typical structures sampled from the space of all possible RNA structures. Our motif frequency study yields another result, that the frequencies of different motifs can be used in machine learning algorithms to classify random and natural RNA with quite high accuracy, especially for longer RNA (eg ROC AUC 0.86 for *L* = 1000). The most important motifs for classification are found to be the number of bulges, loops, and bonds. This finding may be useful in using SS to detect candidates for functional RNA within ‘junk’ DNA regions.

## I. INTRODUCTION

Within evolutionary biology, a long standing debate has centered on whether, and in what ways, development and biases in the genotype-phenotype (GP) map can be a directive force in evolution [1]. In principle, many factors including selection, historical contingency [2, 3], random drift, and biases arising from non-isotropic phenotype variation [4], could have important roles in shaping evolutionary outcomes [5]. Untangling the relative contributions of each factor is difficult hence leading to the protracted debate, and clear data to adjudicate between the various positions has been lacking. Despite these challenges, many studies point to biases in GP maps and development [6–8], and similarly mutation biases [9–11], shaping evolutionary trajectories.

One system which can be used to shed light on the question of bias in evolution is the RNA sequence-to-structure map, where the secondary structure (SS) is taken to be a phenotype describing the pattern of nucleotide base bonding. This map is both computationally and mathematically tractable, for example algorithms exist for predicting SS directly from sequences [12, 13]. At the same time, it is a biologically relevant system, because RNA is a versatile molecule which fulfils many diverse functions in living organisms such as information transfer, catalysis, sensing, and regulation. Moreover, it is well known that RNA SS is important for functional RNA [14–16], and even for messenger RNA [17–19]. SS is also an important determinant of RNA tertiary structure [20]. For these reasons, RNA SS has been studied extensively for many years to elucidate properties of the GP map and as a model system to study evolution [21–28].

It has been known for many years that the RNA sequence-to-structure map is biased in the sense that some SS have disproportionately many sequences underlying them [23]. In other words, there is an exponential variation in the probability of obtaining different SS upon uniform random sampling of RNA sequences, with a small proportion of possible SS accounting for a large fraction of all possible sequences, while many SS only have very few sequences assigned to them.

Dingle et al [29] studied the RNA SS map in the context of the role of bias in evolutionary dynamics. Via computational analysis, they compared natural non-coding RNA of lengths *L* ≤ 126 nucleotides (nt) with randomly generated RNA in terms of the distribution of *neutral set sizes* (ie the number of sequences per SS). They found that the distributions were surprisingly similar. The authors mathematically inferred the distribution of neutral set sizes which would appear from uniform sampling over SS phenotypes (what they called *P-sampling*), and compared it to the observed computationally generated distribution from genotype sampling (what they called *G-sampling*), as well as the distribution from natural data. They also studied other properties of SS again finding similarities for natural and random SS. Moreover, the properties of natural RNA were very different from those of typical SS of the same length within the full space of all possible SS. Their main conclusion was that GP structure appears to strongly constrain which SS are found in nature. Later, Dingle et al [30] extended the earlier study (still using ncRNA with short lengths), by comparing natural and random RNA SS abstracted shapes, not just some of the properties of SS. Their main conclusion was that the shapes of molecules in nature were very similar to shapes derived from random sampling of genotypes, and that the shape frequencies were close to what would be expected from random sampling. The fact that the authors could predict natural shape abundances merely from computer simulations is quite striking. This suggests that the GP map itself is a dominant factor in determining the repertoire and frequency of extant noncoding RNA shapes in nature.

A limitation of these earlier works is that only quite small RNA of lengths *L* ≤ 126 nt were studied, whereas many natural RNA are far larger, leaving the generalisation of the earlier findings to larger — and biologically more interesting — RNA an open question. Much earlier, Fontana et al [21] pioneered comparisons of natural and random RNA finding significant similarities, but their analyses were limited by the data sets available at the time. Here we extend earlier studies by investigating two questions: From the theoretical side, we study much larger non-coding RNA of up to *L* = 3000 nt to determine the role of GP map bias in defining the existing repertoire of RNA, here using structural motifs in addition to RNA shapes.

From the practical side we ask if motif counts can be used to distinguish (or classify) natural vs random RNA, which may be useful in the context of detecting functional RNA in non-coding regions of the genome [31, 32]. This is an important question because while a large fraction of the human genome is transcribed, less than 2% of the genome are protein-coding regions [33]. The function of the other RNA transcripts — or the lack thereof — is a subject of intense research. Furthermore, recent results point to the possibility of uncovering the function of RNA transcripts that have been previously only considered ‘junk’ regions of DNA, with possible functions including stimulation of the immune system during viral infection, in tumor cells or as a consequence of autoimmune disease [34, 35]. Further, with new high-throughput methods to perform full RNA sequencing of a cell, many new RNA transcripts are rapidly being discovered [36]. If functional RNA can be detected with very simple methods such as counting structural motifs, then this would aid in these current research directions.

## II. RESULTS

### A. Abstract RNA shapes

In studies of molecular evolution and bioinformatics, it is common practice to represent RNA SS in dot-bracket form, where a dot represents an unpaired nt base, and a bracket a paired base. Left and right brackets must, therefore, match up; see Figure 1.

**FIG. 1.**
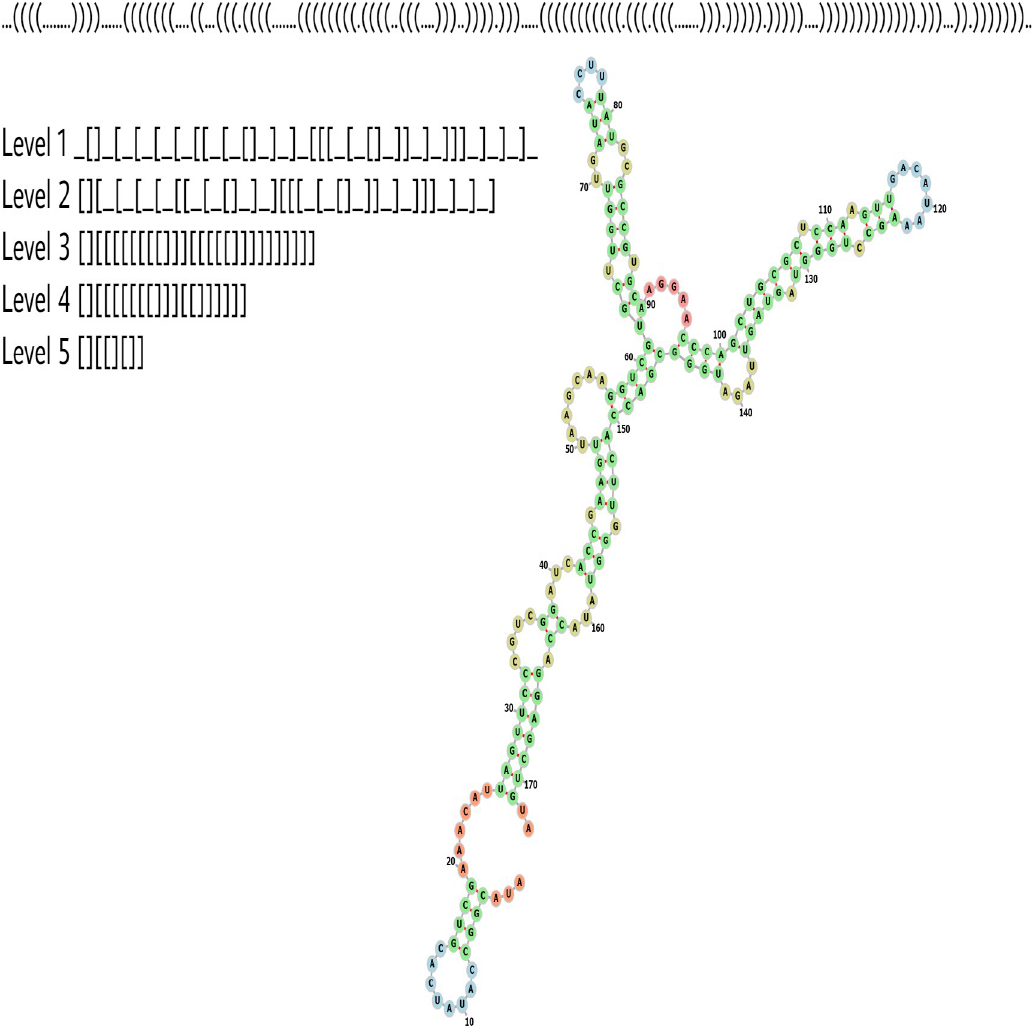
Sepia pharaonis 5S ribosomal RNA, abstract shapes illustration (length is *L* = 173). The dot-bracket and abstracted shapes at higher levels are displayed, corresponding to progressively more coarse grained shapes.

However, studying dot-bracket RNA has some drawbacks: Firstly, a natural RNA such as the Sepia pharaonis 5S ribosomal RNA depicted in Figure 1 may in nature show small variations in length and structural properties, such that an RNA can have slightly different dot-bracket structures. In contrast, the dot-bracket representation defines each different dot-bracket structure as a different RNA phenotype. Hence this type of representation can be seen as perhaps overly detailed for some purposes. Secondly, because of the large variety of different dot-bracket RNA, obtaining statistics about the frequency of a given SS from a database is difficult, because a SS must be found many times to deduce an accurate frequency. This is especially onerous when working with natural databases, which may only have small numbers of samples of each length.

To begin our analysis, following ref. [30] (and see also [37, 38]), we use the RNAshapes [39, 40] method. According to this method, an RNA dot-bracket SS can be abstracted to one of five levels, of increasing abstraction, by ignoring details such as the length of loops, but including broad shape features. Figure 1 illustrates these levels for the Sepia pharaonis 5S ribosomal RNA. The choice of level is a balance between being detailed enough to capture important structural aspects, but not too detailed such that for a given dataset so many shapes are possible, such that very few repeated shapes are found, making it impossible to obtain reliable frequency/probability values. In this current investigation, considering our data sets, we will use level 5 throughout this work.

### B. Nature uses high frequency shapes

To compare shape frequencies for natural and random RNA, we first computationally generate random RNA sequences, then predict their corresponding dot-bracket SS using the popular RNA Vienna package [41]. Then, dot-braket SS are converted into abstract shapes, as described above. To compare to natural SS, we took natural RNA sequences from the RNAcentral [42] database, which is a well-populated database of non-coding RNA, and predict their SS also using the RNA Vienna package. Here we study lengths *L*=100, 200, 300, and 400 for both random and natural sequences. The frequency (or probability) of each phenotype shape *p* is calculated as the fraction of all shapes which have shape *p*. See Methods for more details.

Using the derived frequencies we can make rank plots which have on the *y*-axis the frequency for each shape, and on the *x*-axis the rank of the shape. The highest frequency corresponds to rank 1, the second highest frequency to rank 2, etc. The rank plots are shown in Figure 2, where blue dots indicate random shapes, and in yellow circles are the natural shapes which appeared in the RNA-central database. As is apparent, the shapes in nature tend to have higher frequencies. The number of unique shapes found from sampling 30000 random sequences for each length was 42 (*L*=100), 538 (*L*=200), 3551 (*L*=300), and 12149 (*L*=400). The number of natural sequences were 20223 (*L*=100), 37474 (*L*=200), 19496 (*L*=300), and 34858 (*L*=400). The number of unique resulting natural shapes were 35 (*L*=100), 575 (*L*=200), 2494 (*L*=300), and 8738 (*L*=400). The fraction of unique natural shapes found by random sampling for each length respectively is 34/35 (97%), 397/575 (69%), 1679/2494 (63%), and 4111/8738 (47%). It is interesting that in these cases most of the shapes in nature were found by modest sampling for *L* =100, 200, and 300, and nearly half for *L* = 400, suggesting that nature mainly utilises RNA shapes which are high frequency, and hence easy to ‘find’.

**FIG. 2.**
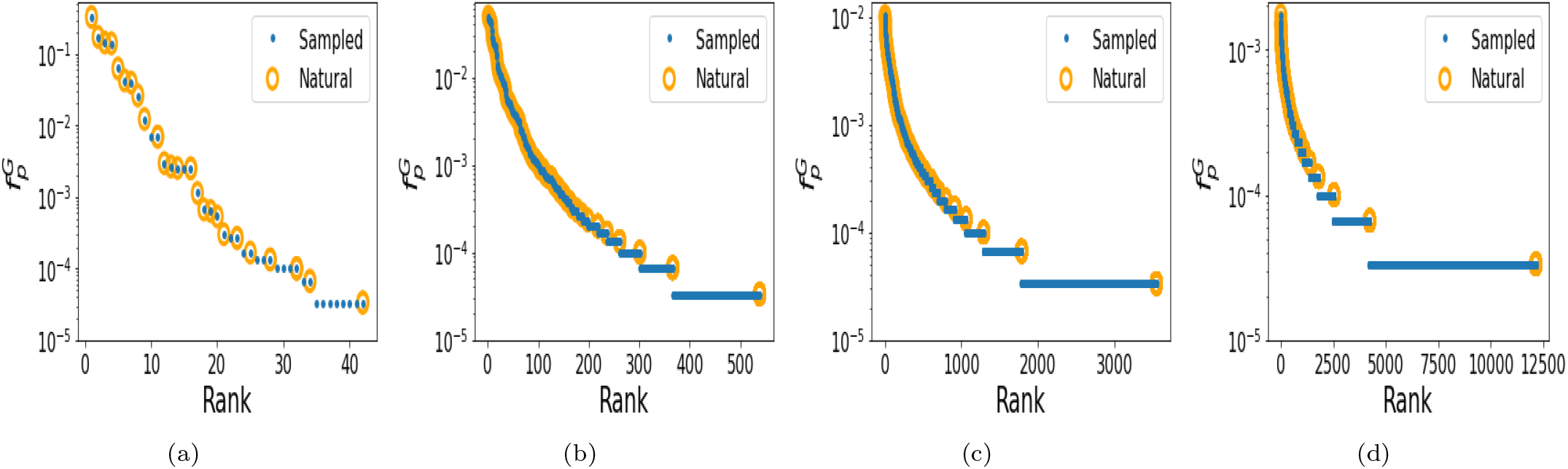
Nature selects frequent structures. Rank plots depicting the frequency 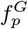 of random (sampled) in blue and natural RNA secondary structure (SS) abstract shapes in yellow. The shapes in nature tend to be those of high frequency. (a) Length *L* = 100 nucleotides (nt); (b) *L* = 200 nt; (c) *L* = 300 nt; (a) *L* = 400 nt.

To help quantify how strong the effect of the phenotype bias is, we can use an estimate of how many shapes actually exist, for each length we have studied. If there are not many possible shapes for a length *L* RNA, then it is not very surprising that modest random sampling finds most of the natural shapes. If there are very many possible shapes, then it is highly unlikely that the relatively modest numbers of natural and random sequences should have shapes that coincide, unless the bias is very strong. Nebel and Scheid [43] made approximate analytic estimates of the number of shapes of length *L* (while ignoring the fact that some shapes may not in fact be designable). For level 5 the number of shapes (*s*_5*L*_) is 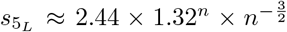 where we have taken the results pertaining to minimum hairpin length of 3, and min ladder length of (which apply to the Vienna folding package). So both are exponential in length *L*. From these equations, we get *s*_5_100__≈10^9^, *s*_5_200__≈10^21^, *s*_5_300__≈10^33^, *s*_5_400__≈10^45^. So we see that the space of possible shapes is astronomical for these lengths. Because only a tiny fraction of these shapes were actually found by sampling, then we can infer that the bias must be very strong for these large RNA, and that nature tends to only use high frequency shapes.

### C. Shape abundance can be predicted from random sampling

Next we compare the frequency 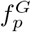 with which RNA shapes appear in random sampling with the frequency *f_p_* with which they appear in the database. This is immediately related to the preceding investigation, but looking at correlations between frequencies is more nuanced than simply looking at whether high frequency shapes appear or not. So, in Figure 3 we show the same data as Figure 2, but now as correlation plots. For *L*=100, 200 and 300 there is a clear positive correlation between the frequency 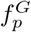 with which shapes appear in randomly generated sequences, and the frequency *f_p_* with which they appear in nature (linear correlations are *r* =0.94, *r* =0.89, *r* =0.79, respectively, all with *p*-values≪ 10^−6^). For *L*=400 the correlation is weak (*r* =0.44, *p*-value≪ 10^−6^), which is likely due to the very noisy frequency data: the total number of possible shapes increases exponentially with length, hence much more data is required to obtain decent statistics for longer lengths. We can conclude that not only does nature typically use highly frequency shapes, but also that the frequency of shapes in nature tends to be similar to that from random sampling. Note that if natural structures were uniformly chosen from the space of possible phenotypes (‘P-sampling’ [29]) then the natural frequencies would be close to uniform, and not correlate with the frequencies of sampled structures.

**FIG. 3.**
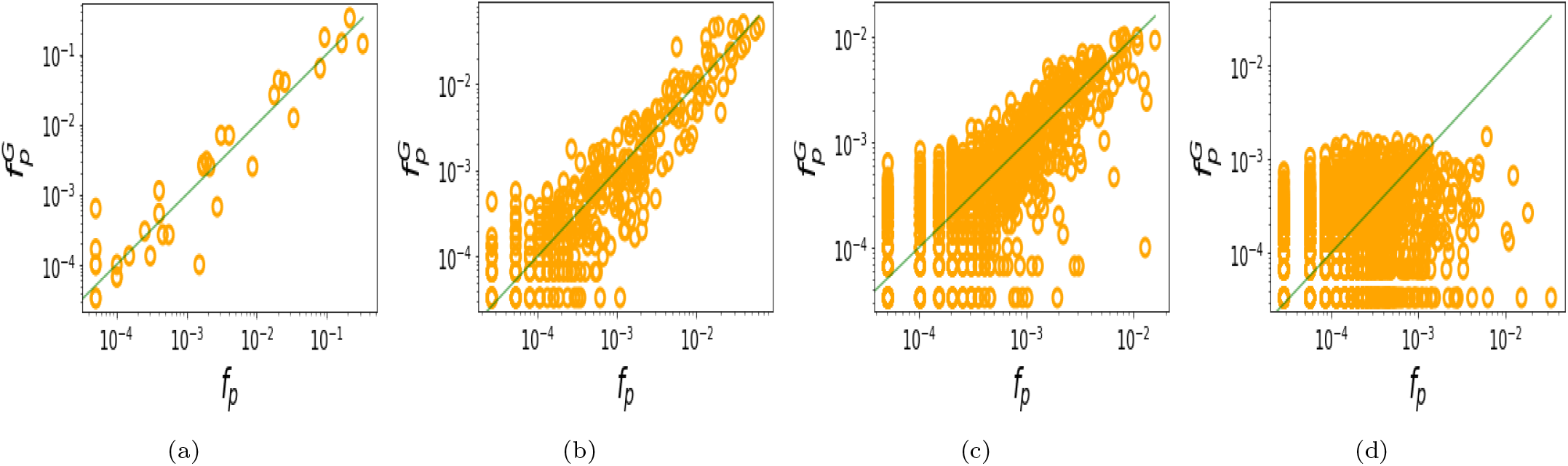
The frequency 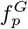 of RNA shapes from random sampling positively correlates with the frequency *f_p_* of shapes in nature. (a) *L*=100, linear correlation *r* = 0.94; (b) *L*=200, *r* = 0.89; (c) *L*=300, *r* = 0.79; (d) *L*=400, *r* = 0.44 (all *p*-values ≪ 10^−6^).

### D. Studying structural motif frequencies for larger RNA

As seen above, for lengths beyond ~300 nt, even at abstract shape level 5 having enough data available to estimate the shape frequencies to high accuracy becomes a challenge. Hence in order to study much larger RNA, we take a different approach: We will compare the computationally folded natural and random structures in terms of fairly easy-to-calculate structural feature motifs, namely the number of helices, bonds, loops, junctions, and bulges. That is, for each RNA dot-bracket SS we will count these motifs and plot them for natural and random RNA with lengths 50 ≤ *L* ≤ 3000 (Methods). In this manner we can see if, for larger RNA, the structural motifs of natural and random SS are similar or not. While the motifs and full RNA shapes are not equivalent, it seems reasonable that if two RNA of the same length have similar numbers of helices, bonds, loops, junctions and bulges, then they are highly likely to have very similar overall shapes.

Figure 4 shows that for each of the 5 motifs, the motif frequency count for natural and random RNA matches quite closely. The bulges and loops appear most different, as inferred from the fact that the lines of best fit for the natural and random data are most divergent, but even then, are still quite similar. The lines of best fit for natural and random RNA for the junctions, helices, and bonds are very close. In the case of the frequency of bonds and frequency of helices, we also plot in Figure 5 an analytic predictions [44] and a computational fit [29], for the expected frequency which would be obtained from P-sampling (ie uniform sampling over all possible SS). As is clear from the figure, neither the natural data nor the random data are similar to the P-sampled values, and in particular the natural and random are far more similar to each other than either is to the P-sampled frequencies.

**FIG. 4.**
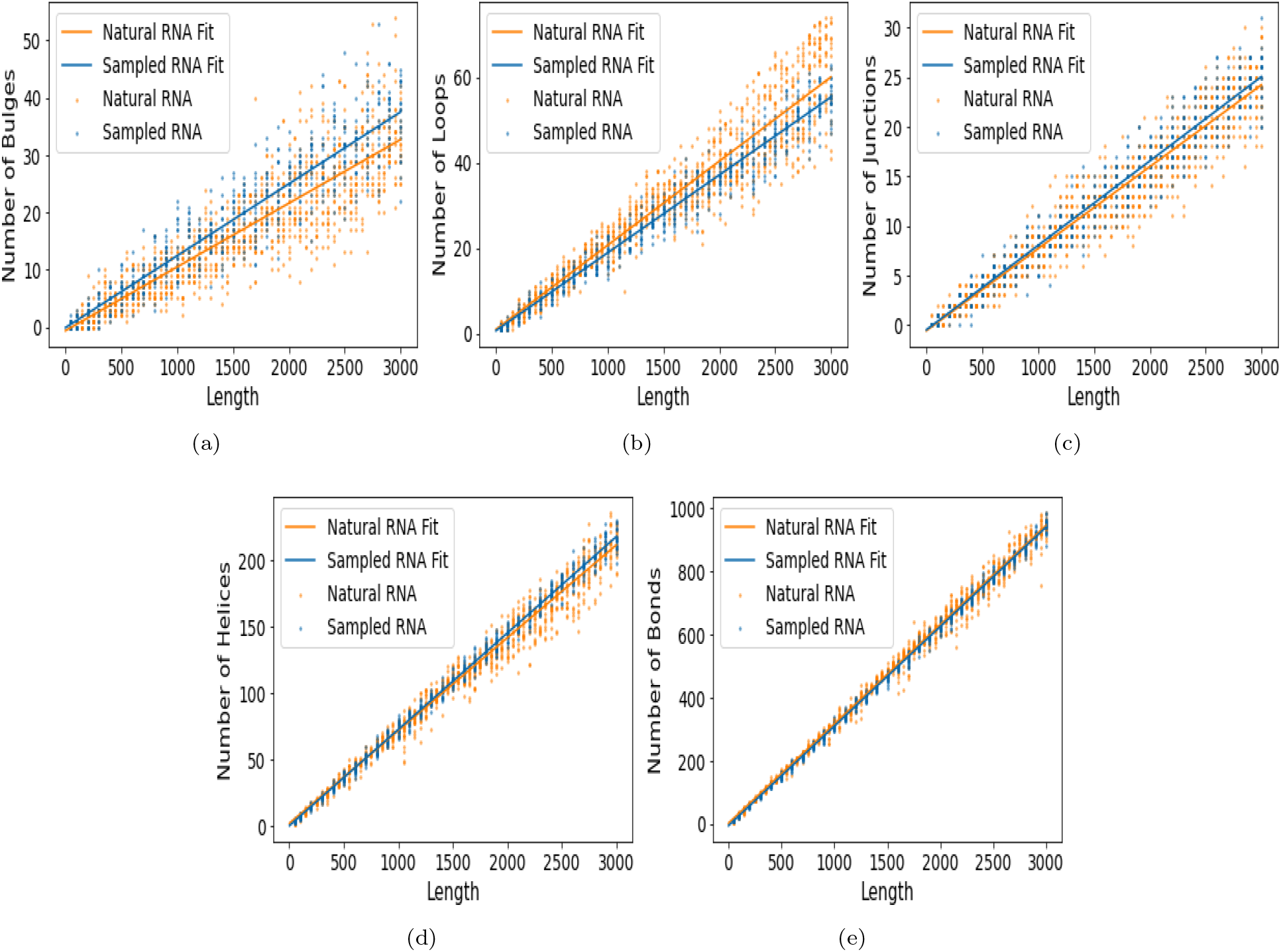
Natural RNA up to *L* = 3000 have similar numbers of bulges, loops, junctions, helices, and bonds compared to randomly sampled RNA. The frequency of bulges and loops appear to differ the most between natural and randomly sampled RNA.

**FIG. 5.**
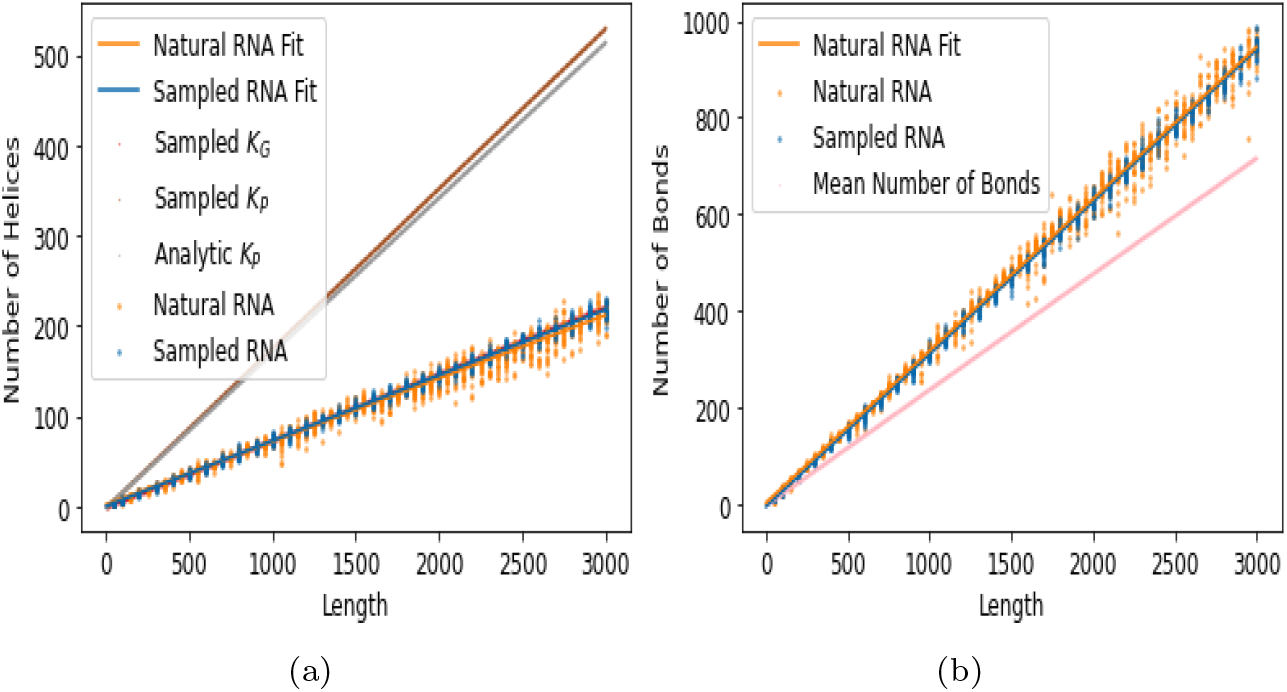
The frequency of (a) helices and (b) bonds observed in natural and random sampling are both very different from those expected from uniform sampling over phenotype SS (P-sampling). *Sampled K_G_* denotes the mean number of helices obtained from random genotype sequence sampling (G-sampling), and *Sampled K_P_* denotes the mean number of helices expected for uniform random sampling over all possible structures (P-sampling) [29]. *Analytic K_P_* denotes the analytic estimated mean number of helices expected for uniform random sampling over all possible structures [44]. Estimated mean number of bonds is also taken from [44].

The linear fits for the motif counts, as a function of length *L*, presented in Figure 4 are given in Table I. The linear fits are *m* = *aL* + *b*, where *m* is the frequency of motif *m* and *a* and b are the slope and intercept of the fit. Table II additionally gives the 95% confidence intervals for the fitting parameters, based on bootstrap sampling.

**TABLE I.** Linear fits *m* = *aL* + *b* for the number *m* of bulges, loops, junctions, helices, and bonds, for natural and random samples.

| Motif | Natural | Random Samples |
| --- | --- | --- |
| Bulges | $0.010L - 0.65$ | $0.013L - 0.12$ |
| Loops | $0.020L + 0.94$ | $0.018L + 0.65$ |
| Junctions | $0.0083L - 0.60$ | $0.0085L - 0.51$ |
| Helices | $0.070L + 1.78$ | $0.073L - 0.049$ |
| Bonds | $0.31L + 2.1$ | $0.32L + 5.8$ |

**TABLE II.** The 95% confidence intervals for the linear fit parameters *a* (slope) and *b* (intercept) given in Table I.

| Motif | Natural | Random Samples |
| --- | --- | --- |
| Bulges | [0.010, 0.011] | [0.012, 0.013] |
| Loops | [0.011, 0.020] | [0.012, 0.018] |
| Junctions | [0.0082, 0.020] | [0.0084, 0.018] |
| Helices | [0.0082, 0.070] | [0.0084, 0.073] |
| Bonds | [0.0082, 0.32] | [0.0085, 0.32] |

| Motif | Natural | Random Samples |
| --- | --- | --- |
| Bulges | [-0.97, -0.35] | [-0.50, 0.28] |
| Loops | [-0.90, 1.2] | [-0.42, 0.86] |
| Junctions | [-0.85, 1.2] | [-0.62, 0.82] |
| Helices | [-0.81, 2.1] | [-0.61, 0.79] |
| Bonds | [-0.79, 2.6] | [-6.4, 0.76] |

As a side comment, in ref. [29] the authors used the neutral network size estimator (NNSE) [45] to compare the neutral set sizes (number of sequences per SS) of natural and random RNA of *L* ≤ 126. On trying to use the NNSE for much larger *L*, we found that the NNSE is not suitable due to increased computational costs and an increasing failure rate in terms of the fraction of sequences for which the algorithm fails to converge.

As has been noted by earlier researchers [46], because the GC content of natural RNA sequences often is biased away from a uniform nucleotide composition value, it is important to check that any observed differences between natural and random RNA are not merely due to a difference in nucleotide composition. Hence we also checked that the observed difference in motif counts persist when using ‘scrambled’ natural RNA sequences (ie randomly permuted natural sequences) instead of uniformly random sequences. The same overall pattern of observations persists, as can be seen in Appendix. This suggests that GC bias alone is not causing the differences between the natural and random samples which we observe.

### E. Biological functions of some high and low frequency shapes

We now look at shapes with high and low frequencies in terms of what biological functions they perform. If an RNA has high frequency, then it can be found by only modest sampling. That is interesting from an evolutionary perspective, because it suggests that natural selection did not have to ‘work hard’ in order to shape the RNA. On the other hand, it would be interesting to see if some natural RNA have very low frequencies, which would suggest that selection had to ‘work hard’ to form that shape. We extracted the highest frequency shape which appeared in random sampling. We then found any molecules in the natural RNA samples from the RNAcentral database which had the same shapes. The most frequent random abstract shapes for the lengths *L* =100, 200, 300, and 400 are [][], [[][][]], [[][][]][] and [][[][][][][]]. In the Supplementary Information, Excel sheets are given which give lists of the RNA names and frequencies of each of frequent shapes. Here we give just the names of the RNA which had the highest frequencies, which were *Sepia pharaonis 5S ribosomal RNA*, *uncultured bacterium partial 16S ribosomal RNA, Hevea brasiliensis miscellaneous RNA*, and *uncultured bacterium bacterial SSU rRNA*, for *L* =100, 200, 300, and 400, respectively.

There is more than one random shape which occurs only once for each length, hence there are numerous ‘least frequent’ shapes in the random samples. Some examples are: the random abstract shape [][[][]][][] with *L* =100 appeared only once in random sampling, and the molecule labelled as *unclassified sequences of pemK RNA* was found to have the same shape. Among the lowest frequency shapes from the random samples for *L* =200 is [[[][]][[][]]][][][], but no natural RNA were found to have this shape. One of lowest frequency shapes from the random sampling for *L* =300 is [][][][[][[][]][][]][][], and this had one occurrence in the natural data, namely *uncultured bacterium partial 16S ribosomal RNA*. For *L* =400, one of the lowest frequency shapes was [[[][][]][[][]]][[][][]][], but this was not found among the natural RNA samples.

## III. CLASSIFYING NATURAL AND RANDOM RNA USING MOTIF COUNTS

### A. Can we use motif frequency to detect functional RNA?

The preceding sections have sought to show that natural and random RNA are overall very similar, especially when compared to the full space of possible RNA SS. However, we have also seen that there are some differences in motif counts between natural and random RNA. Given that we have seen some small differences in the motif frequencies, here we will attempt to distinguish, or classify, natural and random RNA using RNA SS motif counts. One potential application of this would be in detecting functional RNA in supposed ‘junk’ non-coding regions of the genome, as discussed in the Introduction.

Several studies have been undertaken attempting to identify functional RNA or classify different types of RNA. Noteworthy examples include the following. Rivas and Eddy [46] tried to distinguish random and natural RNA, similar to what we consider here. They initially found that SS can be used to distinguish natural and random RNA, but then this ability to distinguish disappeared after adjusting for GC content. Further, they reported that the calculated thermal stability of most fRNA SS are not sufficiently different from the predicted stability of a random sequence to detect functional RNA. Carter et al [47] concluded that using free energy folding values improves function detection beyond just sequence motifs. Bonnet et al [48] showed that microRNA have lower folding free energies than random sequences, but reported that they did not find a good general method to distinguish natural and random RNA, because their method did not work on eg tRNA. Later Washietl et al [49] reported being able to distinguish natural and random RNA using thermal stability of folds. In an effort to classify ncRNAs of different organelle genomes, Wu et al [50] used a machine learning approach involving sequence information and frequency counts of the stem, junctions, hairpin loops, bulge loops, interior loops, and the total loops with more than 3 bases. This work is related to ours in that they used structural motif counts, but differs in that we are not trying to distinguish RNA deriving from different organelles. More recently, Sutanto and Turcotte [51] employed machine learning and structural aspects for the purpose of classifying sequences into specific ncRNA classes. Again this study is related to our current question, but differs in that we are not trying to distinguish different types of natural ncRNA.

### B. Classifying RNA

We now attempt to classify natural and random RNA. The data sets have 5 dimensions, where the features (variables) are the frequency counts of bonds, helices, loops, bulges, and junctions on each SS. There are many algorithms in machine learning which can be used for classification. At first, we use *k*–nearest neighbours (kNN), which is a very common and versatile learning algorithm. We perform 5-fold cross-validation, and classification accuracy is quantified in terms of ROC AUC. Bootstrap sampling is used to obtain 95% confidence intervals for the ROC AUC values. Note that for binary classification an ROC AUC value of ~0.5 indicates very poor classification, no better than guessing classes. Higher values indicate better performance, with 1.0 denoting perfect classification ability.

To experiment we use the following data sets: *L* = 100 with 30000 random and 20223 natural RNA SS, *L* = 400 with 30000 random and 34858 natural RNA SS, and *L* = 1000 with 1000 random and 4836 natural RNA SS. (Note that counting motifs for very large RNA becomes computationally taxing, hence the reduced sample size for *L* = 1000.) The results are presented in Table III, and we see that for longer RNA the classification accuracy is quite high, at around 0.86. The fact that the classification performance is not as high for shorter RNA sequences is expected from Figure 4 in which we saw that the natural and random lines of best fit have slightly different slopes, such that for longer RNA they are more clearly distinguishable. Hence we can expect that classification accuracy is lower for shorter RNA, and higher for longer RNA.

**TABLE III.** ROC AUC values and the 95% confidence interval values for lengths lengths *L* = 100, 400, 1000, and 1000 with adjusted GC content (‘scrambled’ natural RNA). Table (a) uses kNN and (b) uses PLSDA.

| Length ($L$) | Original ROC Area | 95% Confidence Interval |
| --- | --- | --- |
| 100 | 0.72 | [0.72-0.73] |
| 400 | 0.81 | [0.81-0.82] |
| 1000 | 0.86 | [0.85-0.88] |
| 1000 GC adjusted | 0.86 | [0.85-0.87] |

TABLE III. ROC AUC values and the 95% confidence interval values for lengths $L = 100, 400, 1000$ , and $1000$ with adjusted GC content (‘scrambled’ natural RNA). Table (a) uses kNN and (b) uses PLSDA.
| Length ($L$) | Original ROC Area | 95% Confidence Interval |
| --- | --- | --- |
| 100 | 0.70 | [0.70-0.71] |
| 400 | 0.78 | [0.78-0.78] |
| 1000 | 0.83 | [0.82-0.84] |
| 1000 GC adjusted | 0.83 | [0.82-0.84] |

Due to the importance of checking that predictive accuracy is not merely due to a different GC content value [46], for *L* = 1000 we further created a ‘scrambled’ data set, which was made by randomly permuting natural RNA sequences, so as to maintain the same GC content as the natural data. This adjustment for GC content barely lowered the classification performance at all, as shown in Table III.

The kNN method has the benefit of being able to handle arbitrary patterns in data, provided enough data is available. However, it has the drawback that it does not provide an indication of which features (variables) are important in distinguishing the groups. As a different machine learning perspective, we also implemented the partial least squares discriminant analysis (PLSDA) method, which is a linear method that also yields *variable importances*, ie a signed value indicating which features (variables) are most important in distinguishing the groups (larger magnitudes indicate greater importances). The PLSDA method gave ROC AUC values similar but slightly lower than the kNN method, as indicated in Table III.

Figure 6 shows a plot of the variable importances for the natural vs random, and also the natural vs ‘scrambled’ data. The figure indicates the number of bonds is most important for the small RNAs of *L* = 100, the numbers of bonds and loops for *L* = 400, and that the bulges, loops, and bond counts are the most important for *L* = 1000. The number of helices and junctions are relatively unimportant in distinguishing natural and random RNA. Note that in Figure 4 it was visually clear that bulges and loops showed the largest differences between natural and random SS, so it follows that they should appear with large variable importances. Why the bond count also appears as important here even though it does not appear so from looking at Figure 4 is not clear. It may be that there is some kinds of multivariate interaction between the motifs which means that when all variables are considered together, the number of bonds plays an important role.

**FIG. 6.**
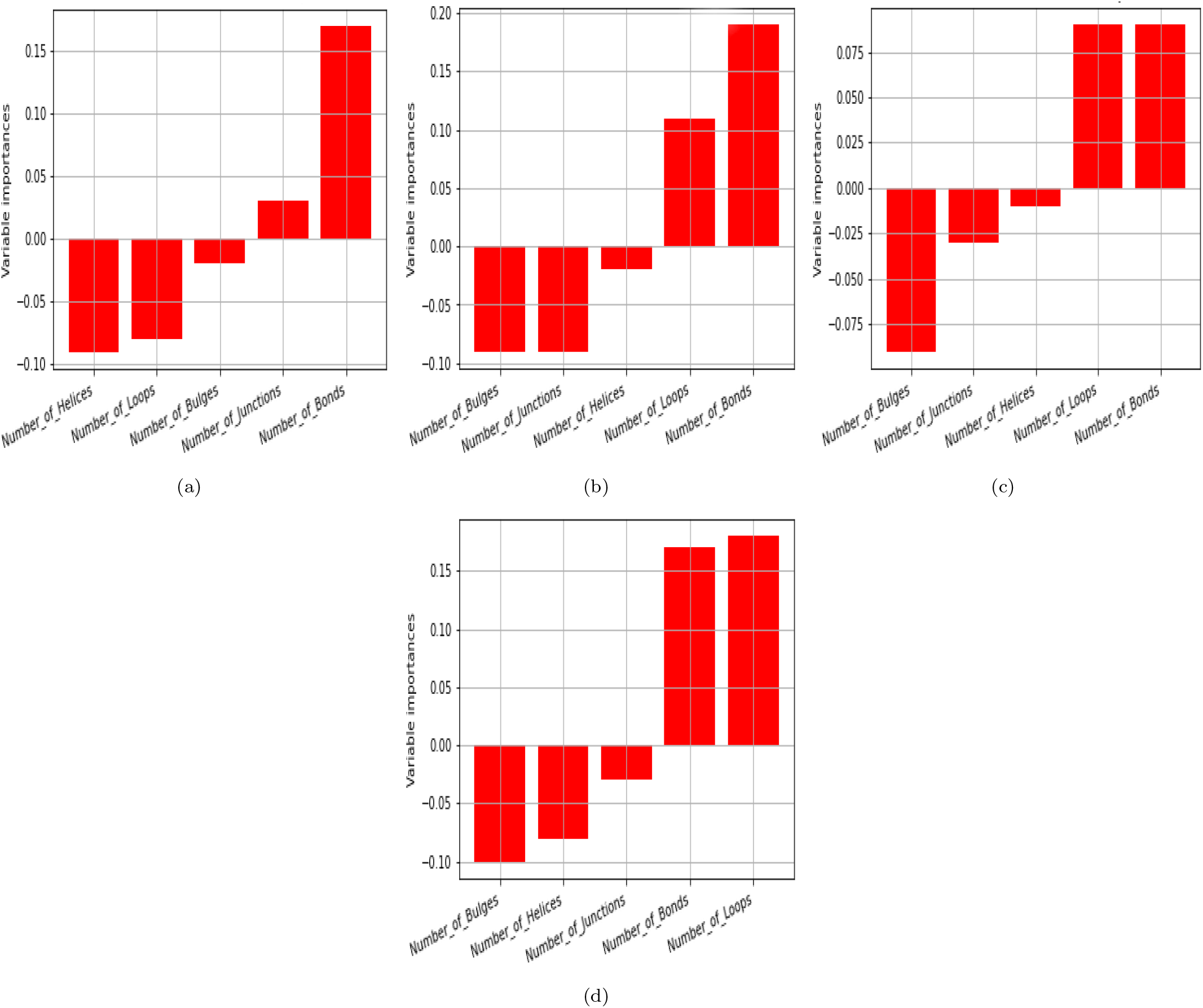
Variable importance plots for different length RNA. (a) Length *L* =100 natural versus random RNA samples, ROC AUC is 0.70 when evaluated using PLSDA (kNN gives 0.72). Bonds are the most important variable. (b) Length *L* =400 natural versus random RNA samples, ROC AUC is 0.78 when evaluated using PLSDA (kNN gives 0.81). Bonds are and loops are the most important variables. (c) *L* =1000, with ROC AUC 0.83 for PLSDA (kNN gives 0.86). Loops, bonds, and bulges are the most important variables. (d) After adjusting the GC content, ROC AUC is 0.83 using PLSDA (kNN, ROC area is 0.86). Loops, bonds, and bulges are the most important variables.

## IV. DISCUSSION

We have compared random and natural non-coding RNA secondary structure (SS), arriving at two main conclusions: Firstly, agreeing with and extending earlier works we showed that natural and random RNA abstract shapes are overall very similar for lengths *L* ≤ 400 nucleotides, and structural motif counts (bulges, loops, helices, junctions, and bonds) are very similar for lengths *L* ≤ 3000. Secondly, despite the overall similarity, we showed that the small differences in motif counts are sufficient for machine learning algorithms to classify natural and random RNA with good accuracy for larger RNA, which may be useful in detecting functional RNA in non-protein coding regions of the genome.

A major motivation for our work was to study the impact of GP map bias on evolutionary trajectories. By “bias” we mean that certain shapes have exponentially more sequences underlying them, as compared to others. Hence random mutations are far more likely to find such preferentially biased shapes, as compared to the others. This bias is a known common property of many GP maps [52, 53]. In this context, adding to earlier works, we suggest that our results here add weight to the case for GP map biases being a substantial, if not actually dominant, player in determining the types of RNA shapes that exist in nature. Put differently, for the larger RNA we have studied here, there are many billions of possible shapes which could appear in nature, but the action of GP map bias restricts the shape repertoire very strongly, leaving natural selection to tune and refine a much smaller set of possibilities. Even in light of earlier works, the overall close similarity of natural and random RNA is rather surprising, because the efficacy of functional RNA is largely determined by their shapes, so *a priori* one would not expect them to be (merely) like random shapes.

Our work accords with experimental studies which have found that diverse structures with potential functionality can be found from samples of random sequences [54, 55], and ref. [56] in which it was shown that natural rRNA have similar structural element properties as compared to random RNA.

Our and earlier findings that natural and random shapes are similar can be explained by the ‘arrival of the frequent’ theory proposed by Schaper and Louis [57], which states that even though selection acts on variation in a population, the GP map biases will strongly shape and constrain which types of variation appear for selection to act on. In their mathematical and computational study, it was shown that even in the presence of natural selection, phenotype bias can still dominate outcomes. See also refs. [27] and [58] for related computational studies and conclusions using RNA and a multi-level GP map. Our results also accord with many other studies which have found that bias can have a strong role in steering evolutionary trajectories [6, 37, 59–64], including the effects of mutation bias [11, 65]. These works support the idea that non-isotropic variation is a significant factor in understanding evolution [66]. Relatedly, it has been shown that the ease of evolutionary accessibility, not relative functionality, can shape which gene network motifs evolve in nature [67].

Another possible explanatory factor for the similarity of natural and random SS is that some fitness-related properties of phenotype shapes are linked to bias. It has been recently argued mathematically that for certain generic fitness requirements based on physics and engineering principles (eg mutational robustness in molecules and efficiency in biological networks), the types of phenotype shapes which have highly optimal values for these generic fitness requirements may also have high probability, ie are favourably biased [68, 69]. In addition to mathematical arguments, a large range of biological examples were presented in support of the theory. Hence there is also a possibility that not only does GP map bias shape the types of variation that is presented to selection, but that there is also some fitness preference for these shapes. Regardless of which of these explanations or combination of explanations holds, it remains an interesting theoretical biology observation that the RNA shapes which appear in nature, and their frequencies, can be predicted by computational and physics-based reasoning.

From a completely different angle, it may be countered that natural selection has adapted RNA SS over time so that the folding rules are tweaked in order to make the types of RNA which are needed by organisms ‘easy’ to generate: rather than the bias shaping which types of RNA are seen in nature, the types of RNA in nature shape the bias. This would explain the similarity of random and natural RNA from a purely selection-based argument. This proposal seems quite unlikely to be valid because RNA folding rules are primarily based on chemistry and physics, so it is hard to see how selection could have had much impact on these rules. Additionally, it has been shown that bias (and probability) are closely related to the information content and complexity of shapes, a general mathematical property [37, 70, 71]. Again information content is not something which selection can substantially alter. See also ref. [72] for a mathematical treatment of neutral set sizes in RNA, again pointing to the fact that bias is related to fundamental mathematical properties of maps, and hence something which is unlikely to be substantially altered by selection.

It is known that a single strand of RNA can fold into more than one possible structure, and some strands even form different structures in vivo and in vitro [73]. Further, even if a given sequence has a minimum free energy SS which dominates over other suboptimal SS, nonetheless the sequence will assume different SS in accordance with a Boltzmann distribution [38, 74]. As is common practice in biology and bioinformatics — as well as the vast majority of earlier RNA SS studies — here we have simplified the GP map by assuming that the minimum free energy SS predicted by the computational folding package is ‘the’ single phenotype. In ref. [30] a brief analysis was made regarding how abstract shapes change if this Boltzmann distribution is incorporated. It was reported that while the dot-bracket SS will fluctuate between various suboptimal folds, the overall shape and hence abstracted shapes do not vary drastically. Hence we do not expect the use of this simplifying assumption to qualitatively affect our conclusions. Nonetheless this simplification forms a limitation of our work.

A limitation of our proposed method to detect functional RNA using SS motif counts is that we implicitly assumed knowledge of the length of the relevant functional RNA sequences, such that we compared, for example, length *L* =1000 natural structures to *L* =1000 random sequences. In practice, given a long non-protein coding region of a genome, we would not know in advance the relevant length to study. Therefore this should be addressed in a future study, before directly applying our classification result in to detecting functional RNA.

Returning to the question of bias, in future work it would be interesting to incorporate different structure prediction methods [75] and especially RNA tertiary structure prediction, if and when it becomes available [76]. Further it would be interesting to study the interplay between bias and selection [63, 77], whether natural or artificial. For example, investigating if the ‘arrival of the frequent’ can be observed experimentally. Possible ways this could be implemented include experimentally via estimating the fitness of RNA molecules as has been done recently for thrombin aptamers [78], or via experimentation combined with deep learning methods to elucidate fitness landscapes, as has been done recently for RNA ligase ribozymes [79]. Such experiments would add data points on which more concrete conclusions could be drawn regarding the role of bias in evolution.

## Supporting information

Excel file with natural RNA names and frequencies

## ACKNOWLEDGMENTS

We thank Ard Louis and Petr Sulc for valuable discussions and suggestions related to this work. We acknowledge financial support from the Kuwait Foundation for the Advancement of Sciences (KFAS) grant number PR19-14SL-02.

## Appendix A: Methods

### 1. Random RNA sequences

We created random sequences using Python, with uniform probability of 0.25 for each nucleotide A, U, C, G. The numbers of random samples were 3 × 10^4^ for *L* = 100, *L* = 200, *L* = 300, *L* = 400 for Figures 2 and 3. For the linear fits to motif counts depicted in Figure 4, the lengths *L* = 50,100,150, 200, 250, …, 3000 were used, and for each length 20 random sequences were made.

### 2. Natural RNA sequences

All natural sequences of lengths *L* = 50,100,150, 200, 250, …, 3000 were downloaded from the RNAcentral database [42] and underwent a cleaning process where any repeated sequence were removed. Any sequences that contained non-standard nucleotide letters were excluded from the samples. For Figures 2 and 3 all acceptable sequences were used. For Figure 4 and the linear fits, because the use of the complete RNAcentral dataset of the specified lengths resulted in a large 4GB file, analysing this dataset would be computationally taxing. Further, because we only required these natural samples for making linear fits, it is not necessary to have such a large dataset to work with. Hence, random sub-sampling was applied on the dataset, where only 20 randomly selected natural sequences of each length were used for the linear fits.

### 3. Folding RNA

We used the popular RNA Vienna package [41] to predict SS from sequences. We set all parameters to their default values (e.g. the temperature *T* = 37°*C*).

### 4. Drawing RNA

Part of Figure 1 was made by drawing an RNA SS using the online tool http://rna.tbi.univie.ac.at/forna/.

### 5. Motif counting

To count the structural motifs, ie the number of bonds, helices, loops, bulges, and junctions on each SS, we used Python code `Secstruc.py` (written by K. Rother). This code is available as a file in the Supplementary Information, along with other code for this project.

### 6. Abstract shapes

Similar to our previous work [30], coarse-grained abstract shapes were used where RNA SS are abstracted from standard dot-bracket notations in different levels. The abstract shapes were obtained using the RNAshapes tool available at bibiserv.cebitec.uni-bielefeld.de/rnashapes and the Bioconda `rnashapes` package available at anaconda.org/bioconda/rnashapes. In order to accommodate the Vienna folded structures, the option to allow single bonded pairs was chosen. In this present work, we used abstract shape level 5, in which no unpaired regions are considered (only if the entire structure is unpaired), and nested helices are combined. Figure 1 in the main text illustrates the abstraction process.

## Appendix B: Adjusting for GC content

In Figure 7 we show linear fits for motif frequencies where instead of purely random sequences, randomly permuted (‘scrambled’) natural sequences are used. As is apparent, adjusting for GC content does not make a substantial difference to motif counts.

**FIG. 7.**
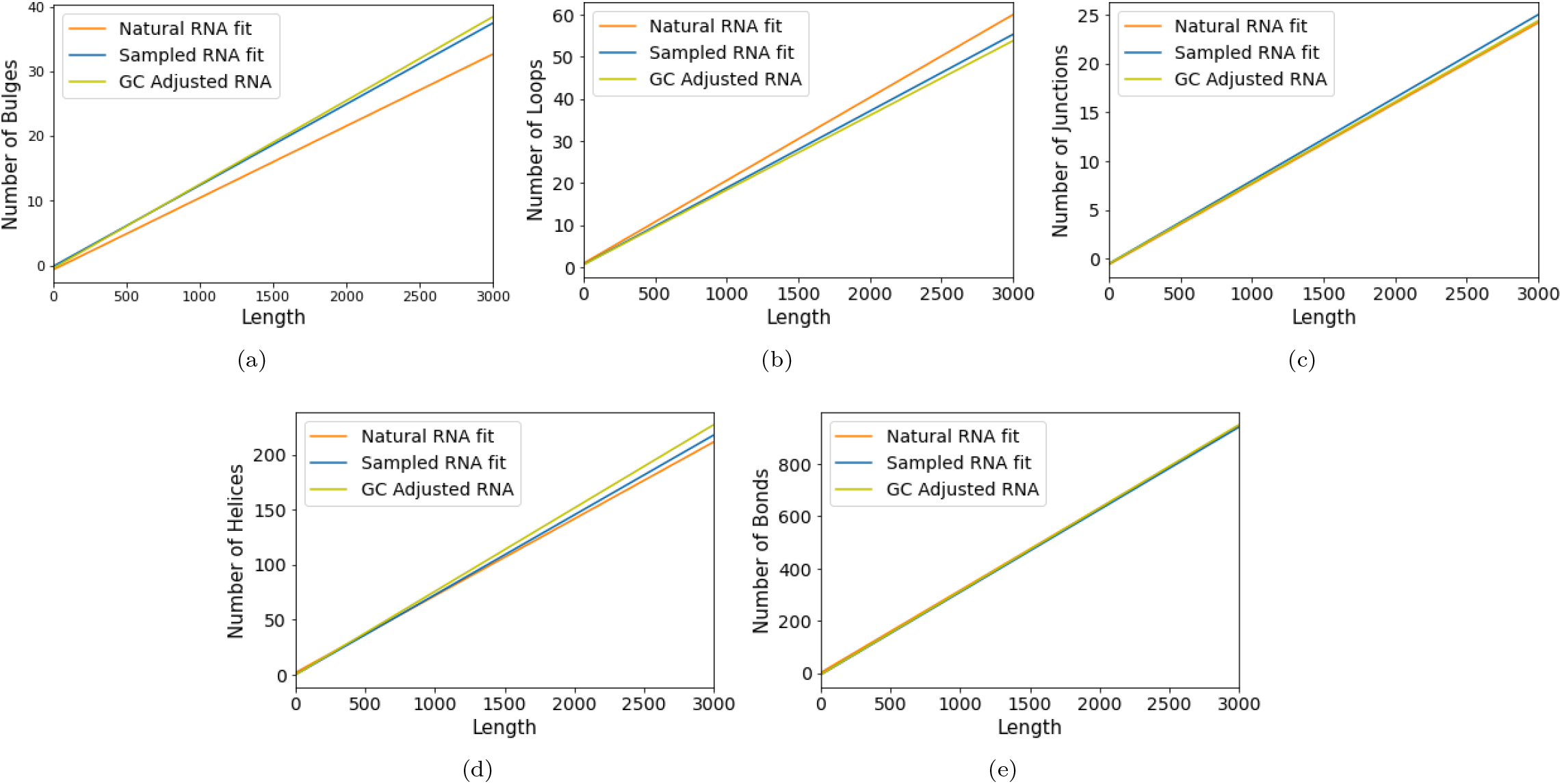
Fits comparing the natural and random RNA, and GC adjusted by scrambling the sequences. GC content does not make a large difference in the scaling of motif counts with *L*, as can be inferred from the fact that the blue and green lines have similar slopes and intercepts.

